# Discrete regulation of β-catenin-mediated transcription governs identity of intestinal epithelial stem cells

**DOI:** 10.1101/2020.05.19.103499

**Authors:** Costanza Borrelli, Tomas Valenta, Kristina Handler, Karelia Vélez, Giulia Moro, Atefeh Lafzi, Laura de Vargas Roditi, George Hausmann, Andreas E Moor, Konrad Basler

## Abstract

The homeostasis of the gut epithelium relies upon continuous renewal and proliferation of crypt-resident intestinal epithelial stem cells (IESCs). Wnt/β-catenin signaling is required for IESC maintenance, however, it remains unclear how this pathway selectively governs the identity and proliferative decisions of IESCs. Here, we demonstrate that C-terminally-recruited transcriptional co-factors of β-catenin act as all-or-nothing regulators of Wnt-target gene expression. Blocking their interactions with β-catenin rapidly induces loss of IESCs and intestinal homeostasis. Conversely, N-terminally recruited co-factors fine-tune β-catenin’s transcriptional output to ensure proper self-renewal and proliferative behaviour of IESCs. Impairment of N-terminal interactions triggers transient hyperproliferation of IESCs, eventually resulting in exhaustion of the self-renewing stem cell pool. IESC mis-differentiation, accompanied by intrinsic and extrinsic stress signalling results in a process resembling aberrant “villisation” of intestinal crypts. Our data suggest that IESC-specific Wnt/β-catenin output requires discrete regulation of transcription by transcriptional co-factors.

## Introduction

Intestinal epithelial stem cells (IESCs) fuel the perpetual replenishment of the epithelial monolayer that lines the gut. They divide on average once a day to give rise to fast cycling progenitors (transit amplifiers, TA), which undergo several rounds of cell division before differentiating into cells of the absorptive (enterocytes) and the secretory lineage (Paneth, goblet, tuft and enteroendocrine cells)^1^. Wnt/β-catenin signalling is required for the maintenance of the intestinal epithelial stem cell pool^2^. However, to date, the gene regulatory networks governing the behaviour of IESCs remain elusive. The activity level of the Wnt pathway alone cannot explain the cell-type specific proliferative rate and self-renewal ability of IESCs, TAs and Paneth cells^3,4^. Hence, discrete regulation of Wnt-target gene expression may determine the identity of the self-renewing pool of IESCs.

A common extracellular signal may be translated into distinct cellular responses, and fate decisions, via the expression of specific cell-autonomous effectors. In the context of Wnt signalling, transcriptional co-factors that interact with β-catenin have been repeatedly implicated in tissue- and cell type-specific modulation of Wnt responses. C-terminal β-catenin interactors, such as CREB-binding protein (CBP) and p300, as well as members of the Mediator and of the RNA polymerase II-associated factor 1 (PAF1) complexes, are promiscuous co-activators that interact, besides with β-catenin, with several transcription factors^5,6^. Conversely, B-cell lymphoma 9 (BCL9) or the paralog BCL9-like (BCL9L) specifically bind an N-terminal moiety of β-catenin, and act as tissue-specific transcriptional effectors^7–11^. While dispensable for normal intestinal homeostasis, BCL9/9L are required for intestinal regeneration upon insults^12–15^, suggesting that they might be necessary for the reconstitution of the stem cell pool. Moreover, compelling evidence has been provided that BCL9/9L plays a crucial role in maintaining tumor stemness in several murine models of colorectal cancer^12–15^.

It remains untested whether distinct regulation of Wnt-signaling outputs is connected to IESC-specific fate determination and self-renewal ability. In this study, we dissect the individual contributions of the C- and N-terminal transcriptional branches of β-catenin to the maintenance of intestinal homeostasis. Using mutant β-catenin alleles that specifically impair the recruitment of N- or C-terminal β-catenin’s co-factors^7^, we show how discrete regulation of Wnt/β-catenin signalling does governs IESC identity and fate. While C-terminal co-actors act as an all-or-nothing switch for Wnt-target gene expression, N-terminal co-activators are responsible for selective transcriptional modulation ensuring proper proliferation and self-renewal of IESCs.

## Results

### Attenuation of N- vs. C-terminal β-catenin transcriptional outputs has contrasting effects on intestinal homeostasis

We previously generated transgenic β-catenin (*Ctnnb1)* alleles harboring mutations that prevent interactions with N- or C-terminal transcriptional co-factors (NTFs and CTFs, respectively)^7^. The D164A mutation abrogates interaction with NTFs, the ΔC truncation abrogates the interaction with the CTFs (Fig. 1a). To overcome embryonic lethality of these alleles, we used compound heterozygous mice carrying one mutant and one conditional β-catenin allele (Extended Data Fig. 1a). Similarly to constitutively hemizygous *Ctnnb1^KO/wt^* mice, *Ctnnb1^D164A/flox^* and *Ctnnb1*^Δ*C*/*flox*^ animals show no overt abnormalities, indicating haplosufficiency of β-catenin for the maintenance of intestinal homeostasis. Combination with the *villin-CreER^T2^* driver^16^ enables the inducible deletion of the conditional β-catenin allele (*Ctnnb1^flox^*) specifically in the intestinal epithelium, thus leaving the Wnt-transcriptional outputs solely under the control of mutant β-catenin (D164A or ΔC). While *villin-CreER^T2^;Ctnnb1^wt/flox^* (control) mice are perfectly viable and indistinguishable from homozygous wild type animals, the presence of only mutant β-catenin is lethal: *villin-CreER^T2^;Ctnnb1*^Δ*C/flox*^ (ΔC) animals exhibit atrophic crypts at 4d post Cre induction (pi) (Extended Data Fig. 1b). This is in accordance with our previous results in *villin-CreER^T2^;Ctnnb1^dm/flox^* animals, which express double mutant (dm) β-catenin harboring both N- and C-terminal mutations^17^. Unlike dm and ΔC animals, *villin-CreER^T2^;Ctnnb1^D164A/flox^* (D164A) animals only reach humane endpoint at 7d pi, as they suffer from severe colitis (Extended Data Fig. 1c). The different intestinal phenotypes that arise upon blocking C- or N-terminal transcriptional outputs of β-catenin are consistent with distinct phenotypic impacts of these mutants during embryonic development^7^.

**Figure 1.**
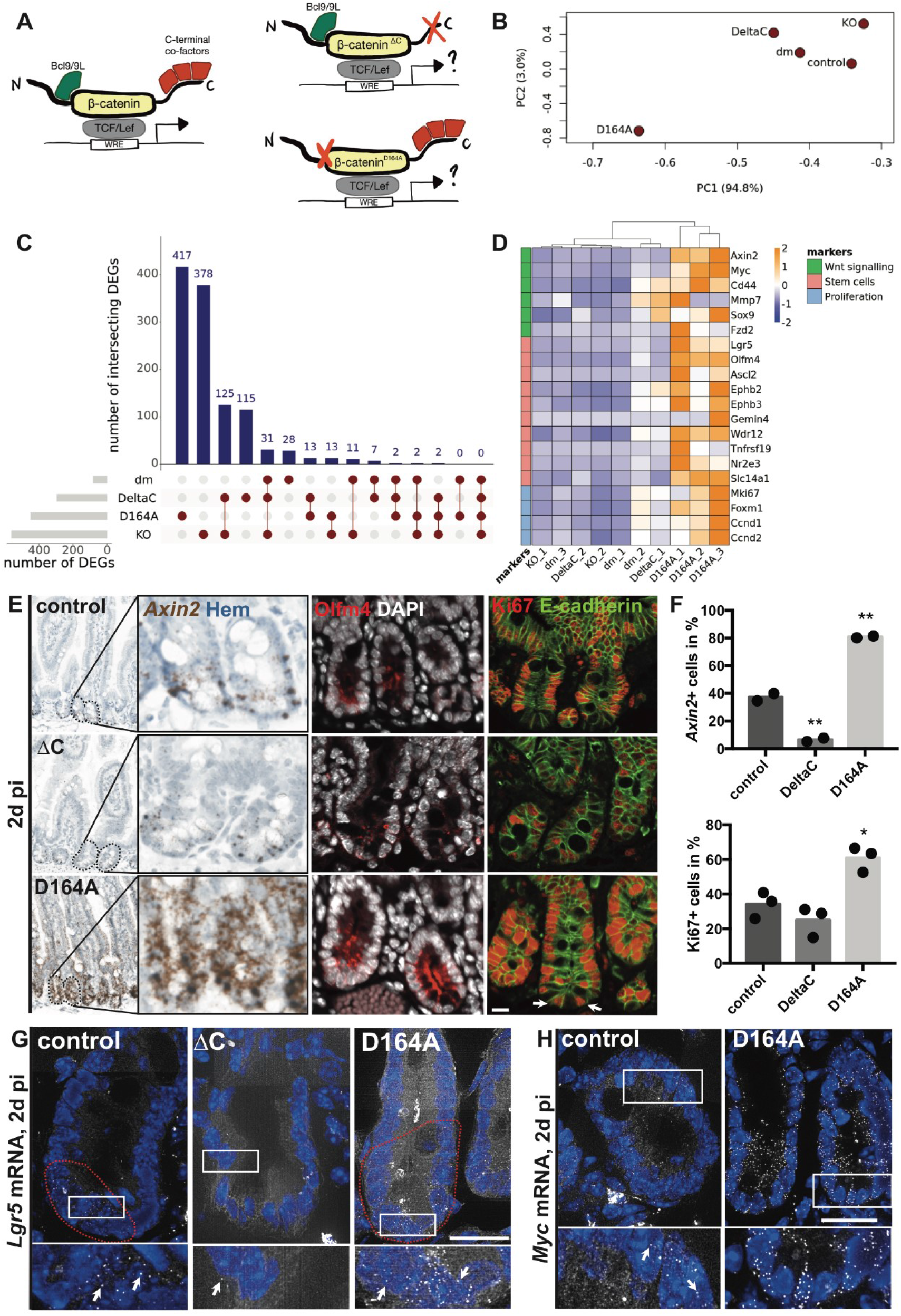
Attenuation of N- vs. C-terminal β-catenin transcriptional outputs has contrasting effects on intestinal homeostasis. A) Scheme of wt and mutant β-catenin proteins with impaired interactions to N- or C-terminal transcriptional co-factors. B) Principal component analysis (PCA) of transcriptome of mutant and control intestinal epithelium 2d pi. PC1 and PC2 explain 94.8% and 3% of the variance, respectively. C) Upset plot of intersecting Differentially Expressed Genes (DEGs) (logFC > |2|, p < 0.01) in ΔC (C-terminal mutant), dm (double mutant), KO (complete loss), D164A (N-terminal mutant), with respect to control. D) Heatmap of normalized FPKM of Wnt and IESC marker genes across hierarchically clustered mutants. Rows are scaled. E) *Axin2* (Wnt-target) mRNA *in situ* hybridization, Olfm4 (IESC marker) and Ki67 (proliferating cells) immunofluorescence of control, D164A and ΔC duodenal crypts 2d pi. Hematoxylin, DAPI (nuclei) or E-cadherin (cell membrane) as counterstain. Arrows indicate Ki67+ cells at crypt bottom. F) Quantification of Axin2+ and Ki67+ cells per crypt. G-H) smFISH of *Lgr5* and *Myc* mRNA. Arrows indicate single mRNA molecules visible as dots. Red dashed line indicates Lgr5+ stem cell compartment. Scale bars, 20 μM. Student’s T-test, unpaired, ** p<0.01, * p<0.05. Abbreviations: KO: *villin-CreER^T2^;Ctnnb1^flox/flox^*, dm: *villin-CreER^T2^;Ctnnb1^dm/flox^* control: *villin-CreER^T2^;Ctnnb1^wt/flox^*, D164A: *villin-CreER^T2^;Ctnnb1^D164A/flox^*, ΔC: *villin-CreER^T2^; Ctnnb1^ΔC/flox^*

Consistent with a fast turnover of intestinal epithelial cells, we observed depletion of wt (flox) β-catenin 2d pi, while the integrity of intestinal crypts remained unaltered (Extended Data Fig. 1d,e). We performed RNA sequencing of bulk preparations of small intestinal epithelium 2d pi from control, ΔC, D164A, dm and KO animals. Principal component analysis indicated surprising differences of impairing N-versus C-terminal interactions on the epithelial transcriptome (Fig. 1b). The differentially expressed genes (DEGs, logFC > |2|, p < 0.05) of ΔC animals broadly overlapped with those of dm and full knock-out (KO) animals, indicating that C-terminally mutated β-catenin is not able to sustain Wnt signalling levels required for proper renewal of the intestinal epithelium (Fig. 1c). Attenuating C-terminal β-catenin transcriptional outputs in the intestinal epithelium caused a loss of IESCs, and an attrition of proliferation in the transient amplifier compartment, as evidenced by IESC gene and proliferative marker expression (Fig. 1d), gene set enrichment analysis (GSEA, Extended Data Fig. 1f)^18^, as well as protein stainings and RNA *in situ* hybridization (Fig. 1e-g). Notably, as previously reported for the β-catenin-dm animals^7,17^, the observed phenotype in ΔC animals is entirely connected to transcriptional outputs, and not attributable to loss of epithelial adhesiveness, as in the case of complete β-catenin loss. Contrary to what was observed in ΔC mice, the transcriptomic changes induced in β-catenin-D164A animals (i.e. N-terminal mutant) only minimally overlapped with those induced by the loss of β-catenin (Fig. 1c). The exclusivity of DEGs in D164A-mutant can be partially attributed to a D164A-specific enrichment of genes expressed by infiltrating immune cells (Extended Data Fig. 1g). As opposed to what we observed in ΔC crypts, the expression of Wnt targets, IESC genes and proliferation markers, was significantly increased in D164A crypts 2d pi (Fig. 1e-f). Moreover, D164A mutant crypts displayed a significant increase of the *Lgr5*+ area, indicating an expansion of the IESC compartment (Fig. 1g, Extended Data 1h). Furthermore, the presence of Ki67+ cells at the bottom of D164A crypts indicates increased IESC proliferation (Fig. 1e,f). Consistent with hyperproliferation in the stem cell compartment, *Myc* expression was upregulated in the bottom of the crypt, as shown by single molecule fluorescent *in situ* hybridization (smFISH) (Fig. 1h). These results indicate that attenuating β-catenin’s N-terminal transcriptional outputs increased proliferation of IESCs and resulted in an expansion of the stem cell compartment. On the contrary, preventing interactions to CTFs completely represses the Wnt-outputs, including proliferation and IESC-associated genes, hence compromising stem cell maintenance.

### Intestinal crypts lacking the output via of N-terminal recruited co-factors exhibit stem cell loss and secretory hyperplasia 4d pi

Unexpectedly, in contrast to 2d pi, expression of Wnt-target genes, as well as IESC genes and proliferative markers was completely abrogated in D164A crypts 4d pi (Fig. 2a). Moreover, crypt morphology was profoundly altered by the appearance of large granular cells that express the secretory lineage marker Sox9 (Fig. 2a) and are double positive for lysozyme (Lyz) and mucins (stained by alcian blue, AB) (Fig. 2a,b). These so-called intermediate cells, which share both Paneth (lysozyme) and goblet cell traits (mucins), are the result of stem cell mis-differentiation, and have been observed upon experimental perturbation of Notch, Wnt or EGF signalling in the intestinal epithelium^19–23^. However, the molecular mechanisms underlying their appearance in the tissue is largely unknown. Interestingly, a shift towards secretory lineage differentiation was observed upon deletion of N-terminal co-factors BCL9/9L in APC^min^ tumors^15^, suggesting that the NTFs might repress specification of the secretory lineage.

**Figure 2.**
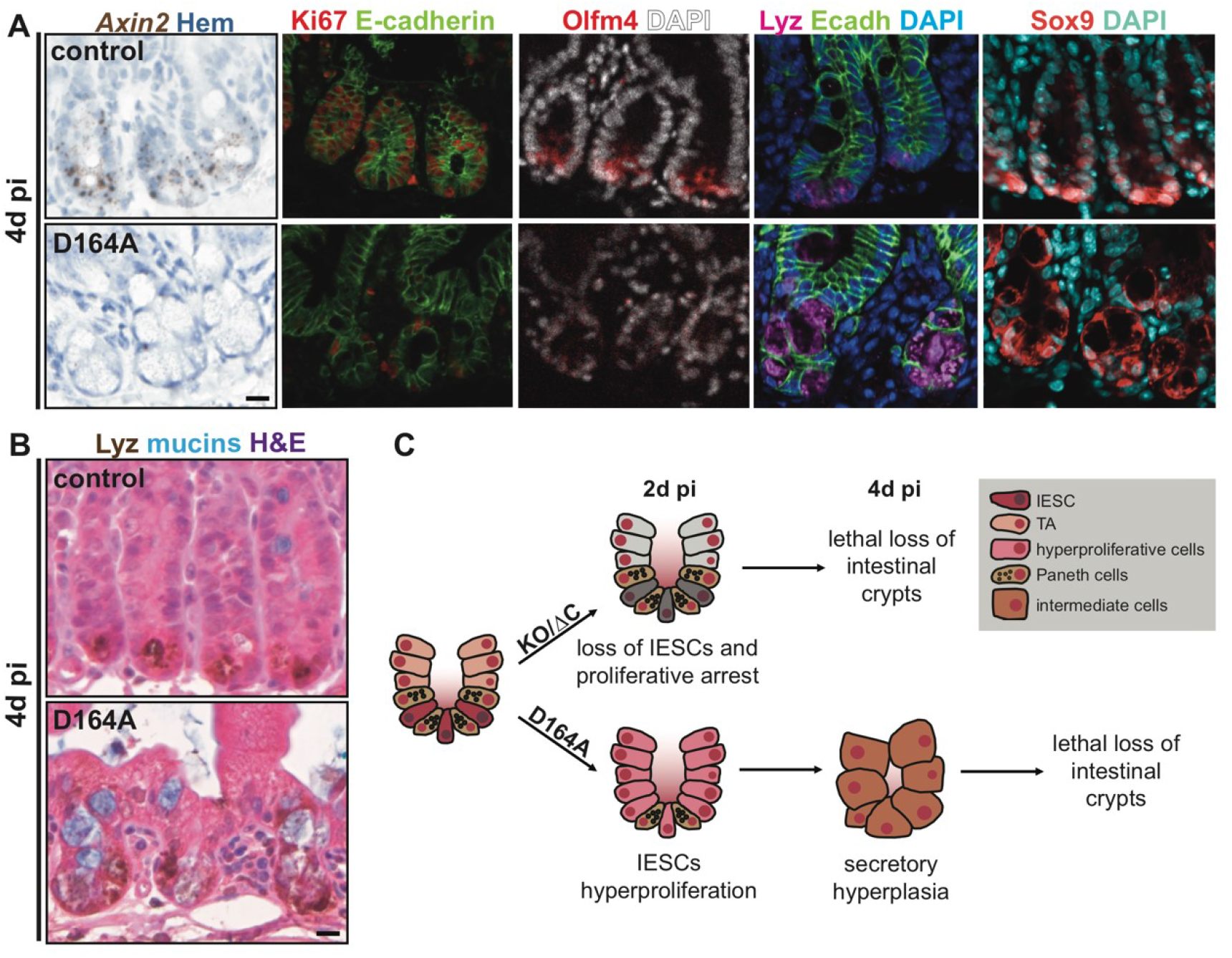
Hyperproliferative intestinal crypts lacking the output via of N-terminal recruited co-factors exhibit stem cell loss and secretory hyperplasia 4d pi. A) *Axin2* (Wnt-target) *in situ* hybridization, Olfm4 (IESC marker), Ki67 (proliferating cells), lysozyme (Lyz, Paneth cells) and Sox9 (Paneth cells per-cursors) immunofluorescence of control and D164A crypts 4d pi. Hematoxylin, DAPI (nuclei) or E-cadherin (cell shape) for counterstain. B) Lyz-HRP (Paneth cells) and alcian blue-stained mucing (goblet cells) double staining. Hematoxylin-eosin as counterstain. C) Working model summarizing impacts of missing contribution of N- vs. C-terminal β-catenin co-factors.

Taken together, these results indicate that neither C- nor N-terminally mutated β-catenin is haplosufficient in the intestinal epithelium, as these mutants are not able to substitute for the wild type allele. However the cellular dynamics leading to crypt collapse are profoundly distinct (Fig. 2c). Similar to the complete deletion of β-catenin, blocking its CTF recruitment rapidly caused a lethal loss of stem cells. On the other hand, impairing interaction with BCL9/9L triggered transient hyperproliferation of IESCs, followed by their aberrant differentiation.

### Hyperproliferative IESCs expressing only *Ctnnb^1D164A^* have an increased ability to form organoids

Recent work indicated that intestinal organoids could not be established from BCL9/9L-deficient crypts isolated 4d pi^14^. However, we hypothesized that expansion of the proliferative IESC compartment in D164A crypts observed 2d pi would translate into increased organoid formation rate. Thus, we devised an organoid formation assay to test this (Fig. 3a): we isolated intestinal crypts from *villin-CreER^T2^;Ctnnb1^wt/flox^* (control) and *villin-CreER^T2^;Ctnnb1^D164A/flox^* (D164A) animals 0, 2 and 4d post Cre induction, seeded equal amounts in Matrigel and quantified the number of organoids formed 7d post seeding. Intriguingly, hemizygous control crypts showed a drastic reduction in organoid formation rate upon loss of the conditional allele (Fig. 3b). Indeed, in contrast to the *in vivo* situation, we found that β-catenin is not haplosufficient *in vitro*: neither crypts isolated from constitutively hemizygous mice, nor upon acute Cre induction (*in vivo* and *in vitro*) can be propagated as intestinal organoids (Fig. 3c and Extended Data Fig. 2a). Moreover, crypts isolated from *Ctnnb1*^Δ*C/flox*^ and *Ctnnb1^dm/flox^* animals also fail to grow *in vitro* (Extended Data Fig. 2b, indicating that two copies of C-terminally transcriptionally active β-catenin alleles are necessary to maintain Wnt-target gene expression required by intestinal organoids. The differential Wnt/β-catenin-signalling requirements of intestinal crypts *in vitro* might be due to the fact that organoids lack a mesenchymal support niche, and that they closely resemble a regenerative, rather than homeostatic state of intestinal crypts. Nonetheless, D164A crypts isolated 2d pi exhibited a 1.6-fold increase in organoid formation rate compared to non-induced (*Ctnnb1^D164A/flox^*) crypts, and a 10-fold increase compared to the respective control (Fig. 3c). Morphologically, organoids derived from hyperproliferative crypts showed increased cell proliferation and budding (Fig. 3c). Consistent with the transient nature of the hyperproliferation observed *in vivo,* these organoid lines obtained from D164A crypts 2d pi cannot be maintained in culture, and died upon first passaging (post splitting panels, Fig. 3c). Crypts isolated from D164A animals 4d pi failed to establish organoids *in vitro,* in agreement with the loss of stem cell and proliferation markers observed in the tissue (Fig. 3b). In sum, these results *in vitro* recapitulate and functionally underscore our observations *in vivo*.

**Figure 3.**
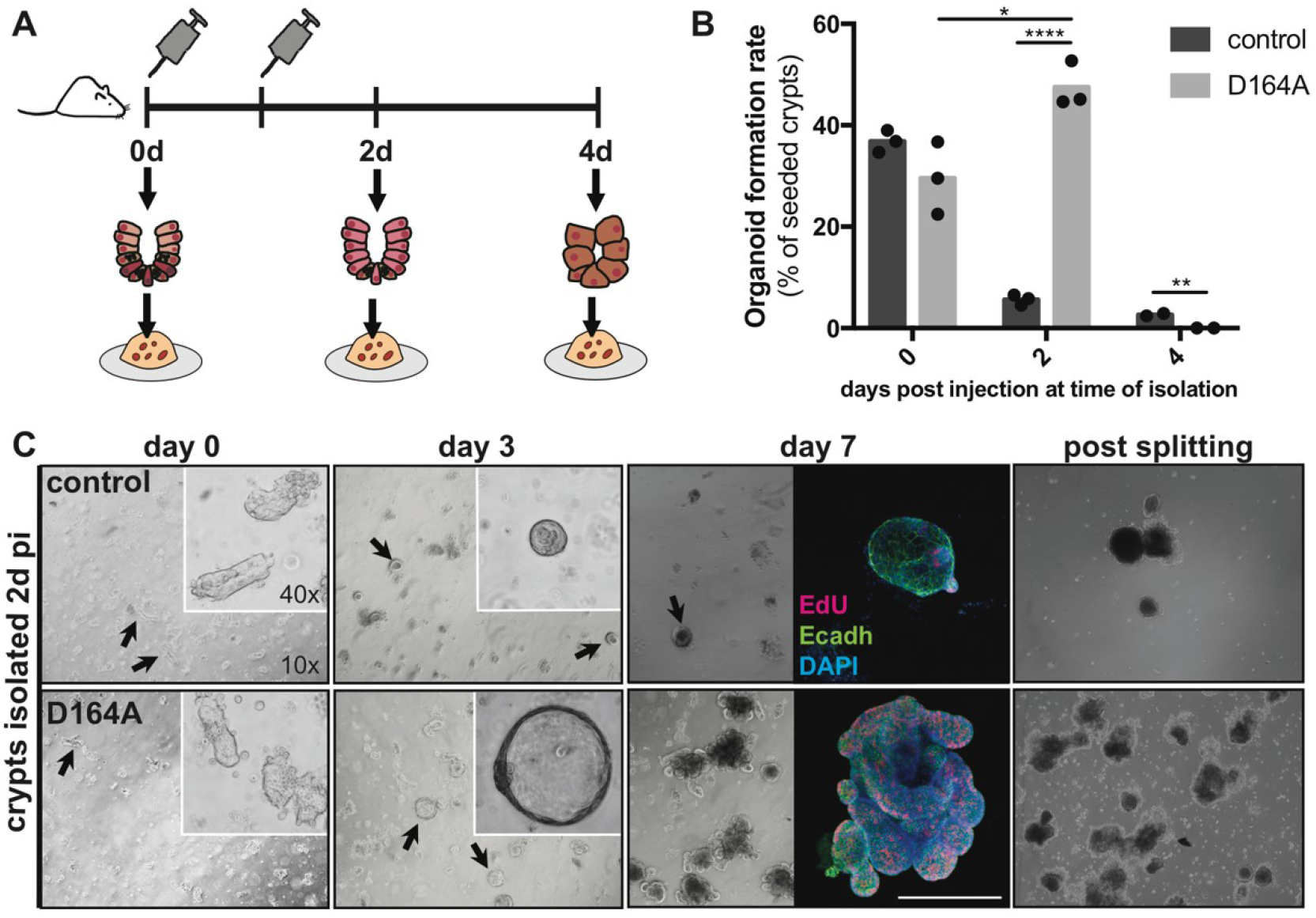
Hyperproliferative IESCs expressing only *Ctnnb1*^*D164A*^ have an increased ability to form organoids. A) Experimental workflow of organoid formation assay. B) Organoid formation efficiency (in %) of control (*villin-CreER^T2^;Ctnnb1^wt/flox^*) and mutant (*villin-CreER^T2^;Ctnnb1^D164A/flox^*) crypts isolated 0, 2 and 4d pi. C) Brightfield images of control and mutant organoids formed from crypts isolated 2d pi. Inset at day 7 EdU+ shows EdU staining (proliferating cells). E-cadherin (cell shape) and DAPI (nuclei) as counterstain. Scale bars, 20 μM. Student’s T-test, unpaired, **** p < 0.0001, ** p < 0.01, * p < 0.05 *.

### Myc-E2F-driven proliferation leads to exhaustion of the stem cell pool in N-terminal mutant (D164A) animals

To dissect the gene regulatory networks underlying the transient hyperproliferation of IESCs triggered by impaired β-catenin-NTF interactions, we performed droplet-based single cell RNA sequencing (scRNAseq) of small intestinal crypts isolated from control and D164A intestines at 0, 2 and 4d pi. We focused our analysis on the gene expression changes occurring over time in stem cells and early progenitors (SCEP cells), which we annotated manually based on marker gene expression (Extended Data Fig. 3a-c). To disentangle the transcriptomic effects caused by impaired N-terminal interactions from those arising due to the acute loss of the conditional β-catenin allele, we devised a normalization procedure using the control time series data (detailas in Methods).

Confirming the data presented above (Fig. 1d-g), D164A SCEP cells exhibited transient upregulation of IESC and proliferation markers 2d pi, followed by increased expression of differentiation markers 4d pi (Extended Data Fig. 3d). This is highlighted by diffusion maps^24^ as a shift along the stem cell differentiation continuum: SCEP cells isolated 2d pi have a significantly more ‘‘stem-like’’ distribution, whilst SCEP cells from D164A crypts 4d pi are more shifted towards differentiation (Fig. 4a,b). The upregulation of stem cell markers 2d pi was accompanied by a significant increase in the mean expression levels of cell cycle-related genes^25^ (Fig. 4c). Supervised pseudotime analysis^26^ confirmed that stem cell and proliferation markers exhibit a similar expression profile over time, increasing 2d pi, and subsequently dropping 4d pi (Fig. 4d). We had observed increased *Myc* expression in D164A crypts 2d pi (Fig. 1h), we therefore asked if Myc-driven proliferation was causing increased cell cycle rate of SCEP cells. At 2d pi, Myc-dependent Wnt-targets were significantly enriched among DEGs between 0d and 2d pi (MSigDB name: M1757, normalized enrichment score (NES)=1.68, p<0.05, from^27^) (labeled as APC-targets in Fig. 4e). Moreover, the signature of E2F - the major cell cycle regulator, and a Myc-target - was significantly enriched (M5901, NES=1.56, p<0.05). Also enriched were genes involved in cell cycle (M14460, NES=1.65, p<0.05) and G2M checkpoint (M5901, NES=1.72, p<0.05) (Fig. 4d). These proliferation signatures were all significantly depleted 4d pi (NES<−1, p<0.05), reflecting abrupt proliferative arrest (Fig. 4c,d). We wondered whether the observed Myc-E2F-induced proliferation burst induced an en-bloc conversion of D164A IESCs to TAs, which are known to exhibit higher expression of proliferative Wnt targets^28^. To test this hypothesis, we trained a logistic regression model on TAs and IESCs obtained from a publicly available single cell dataset of murine small intestinal epithelium^29^, and tested SCEP cells from 0d and 2d pi against this model. Comparison of the distribution of predicted model responses from the two timepoints shows a significant shift of 2d pi cells towards TA traits (Extended Data Fig. 3d). Cumulatively, our results indicate that impairing β-catenin-NTF interactions transiently activates an Myc-E2F-driven proliferation program in IESCs, which results in exhaustion of the self-renewing IESC pool.

**Figure 4.**
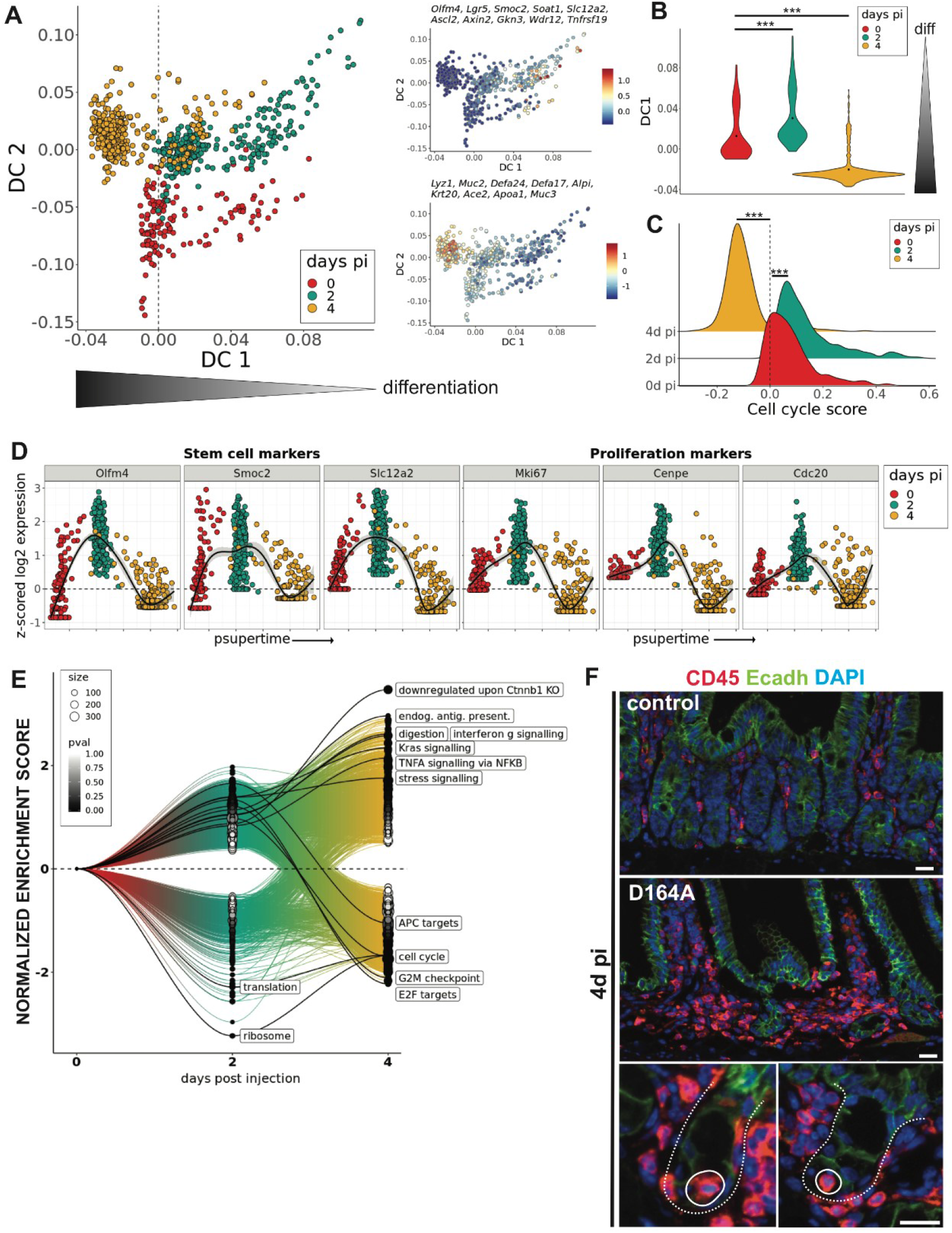
Myc-E2F-driven proliferation is followed by stress signalling and loss of stem cells in N-terminal mutant (D164A) crypts. A) Diffusion map for normalized D164A stem cells and early progenitors (SCEPs). Cells (points) colored by timepoint after Cre induction (days pi). Diffusion component 1 (DC1) represents differentiation pseudotime. On the right, cells colored by their average expression of stem cell (top) and differentiation markers (bottom). B) Distribution of DC1 score in D164A SCEP cells over time (***p < 0.001, Wilcoxon test). C) Distribution of cell cycle score (average expression of cycling genes) in SCEPs over time (***p < 0.001, Wilcoxon test). C) Expression profile of stem cell and proliferation markers in SCEPs ordered along supervised pseudotime (psupertime). Cells colored by timepoint. D) Gene set expression analysis GSEA on time-varying genes in SCEPs across timepoints. Normalized enrichment score (y-axis) over time (x-axis) of MSigDB gene sets. Dot size indicates size of enriched gene set. Dot color indicates p value of enrichment. Selected gene sets labeled (identifiers in text). E) CD45 immunofluorescence in control and mutant 4d pi. Details show immune cell (circle) infiltration within mutant crypts (dashed line). Scale bars, 20 μM.

### Environmental and cell-intrinsic stress signals induce mis-differentiation

Gene set enrichment analysis (GSEA) of time-varying genes in SCEP cells also revealed an increasing extrinsic and intrinsic stress signaling in N-terminally mutated (D164A) crypts. Signatures of TNFα signalling and interferon-γ signalling are significantly upregulated in D164A SCEP cells (M5913, NES=2.32, p<0.001 and M5890, NES=2.08, p<0.001, respectively). Accordingly, the cytokine-induced genes *Irf7*, *Fos*, *Jun*, and *Junb* are among the most upregulated transcription factors according to our supervised pseudotime analysis (Extended Data Fig. 3f). Moreover, we found genes involved in endogenous antigen presentation to be significantly upregulated over time in D164A SCEP cells (M16750, NES=3.01, p<0.001). Interestingly, a proliferative subset of IESCs has recently been described to function as unconventional antigen presenting cells, orchestrating interactions with immune cells in the intestine via expression of MHC class II complexes^30^. In line with these results, and together with the lymphocyte recruitment indicated by the RNASeq data (Extended Data Fig. 1g), we observed infiltration of CD45+ cell in the pericryptic zone, as well as within crypts, of D164A animals 4d pi (Fig. 4f).

The unfolded protein response (UPR) is a major cell-intrinsic stress response. Key mediators of the UPR - *Xbp1*, *Creb3, and Creb3l3 -* were among the upregulated transcription factors over time (Extended Data Fig. 3f). *Creb3l3* is normally a strongly zonated gene; its expression increasing along the crypt-villus axis, with peak expression at the villus tip^31^. We confirmed its ectopic expression in D164A crypts 4d pi using smFISH (Extended Data Fig. 3g). Consistent with an UPR, genes involved in translation initiation and ribosomal function are significantly downregulated in D164A crypts (respectively M27686, NES=-2.52, p<0.05 and M189, NES=-3.26, p<0.001) (Fig. 4e). Interestingly, both UPR^32^ and interferon-γ signalling^30^ have been shown to induce differentiation of IESCs. Furthermore, antigen presentation is intrinsically linked to UPR^33^. For instance, prior to the discovery of its role in UPR, Xbp1 was originally discovered as a regulator of class II major histocompatibility genes^34,35^. Altogether, these results suggest that, following hyperproliferation of IESCs, a combination of environmental (cytokines) and cell-intrinsic stress induces loss of stem cells and secretory hyperplasia of D164A crypts, resulting in loss of stem cells and subsequently failure of epithelial homeostasis.

### The chromatin landscape of D164A crypts reflects the transition to proliferative arrest and mis-differentiation

Our analyses revealed rapid and profound transcriptomic and morphological changes in the crypts upon impairment of β-catenin-NTF interactions (D164A mutant). We reasoned that profiling the chromatin landscape of D164A crypts would provide insight into the gene regulatory mechanisms responsible for such drastic reprogramming of epithelial cells. To this end, we performed Assay for Transposase-Accessible Chromatin using sequencing (ATACSeq) of control and D164A crypts isolated 2d pi. Differential peak analysis revealed that 1469 ATAC-peaks are differentially accessible (logFC > |1| & p < 0.01) in D164A crypts, compared to control crypts (Fig. 5a,b). We subjected these peaks to motif analysis with HOMER^36^, and found a significant enrichment (Benjamini-corrected q-value < 0.001) of motifs associated with transcription factors involved in intestinal differentiation, such as hepatocyte nuclear factors (HNF4a,1,1b), ETS-related transcription factors (EHF, ELF3,5) Caudal Type Homeobox (Cdx2,4), GATA-binding factors and Krüppel-like factors (Extended Data Fig. 4a). These findings correlate with increased expression of secretory and absorptive lineage markers in D164A crypts seen at 4d pi. Moreover, peaks annotated to Wnt target genes (*Tcf7l1/2, Axin2, Fzd7, Sp5*), Wnt-inhibitors (*Tle3, Trabd2b, Wif1, Dab2ip, Amer3*) and cell cycle genes (*E2f1, Cenpf, Cebpe*) Fig. among the lost peaks (Fig. 5b). These results indicate that the chromatin landscape 2d pi preludes the proliferative arrest and secretory hyperplasia observed in N-terminal mutants (D164A) 4d pi. Indeed, we found that a significant fraction of pathways overrepresented in D164A vs control crypts according to our ATAC-peak analysis (2d pi) were also overrepresented in the transcriptome of D164A mutant crypts 4d pi (Extended Data Fig. 4b).

**Figure 5.**
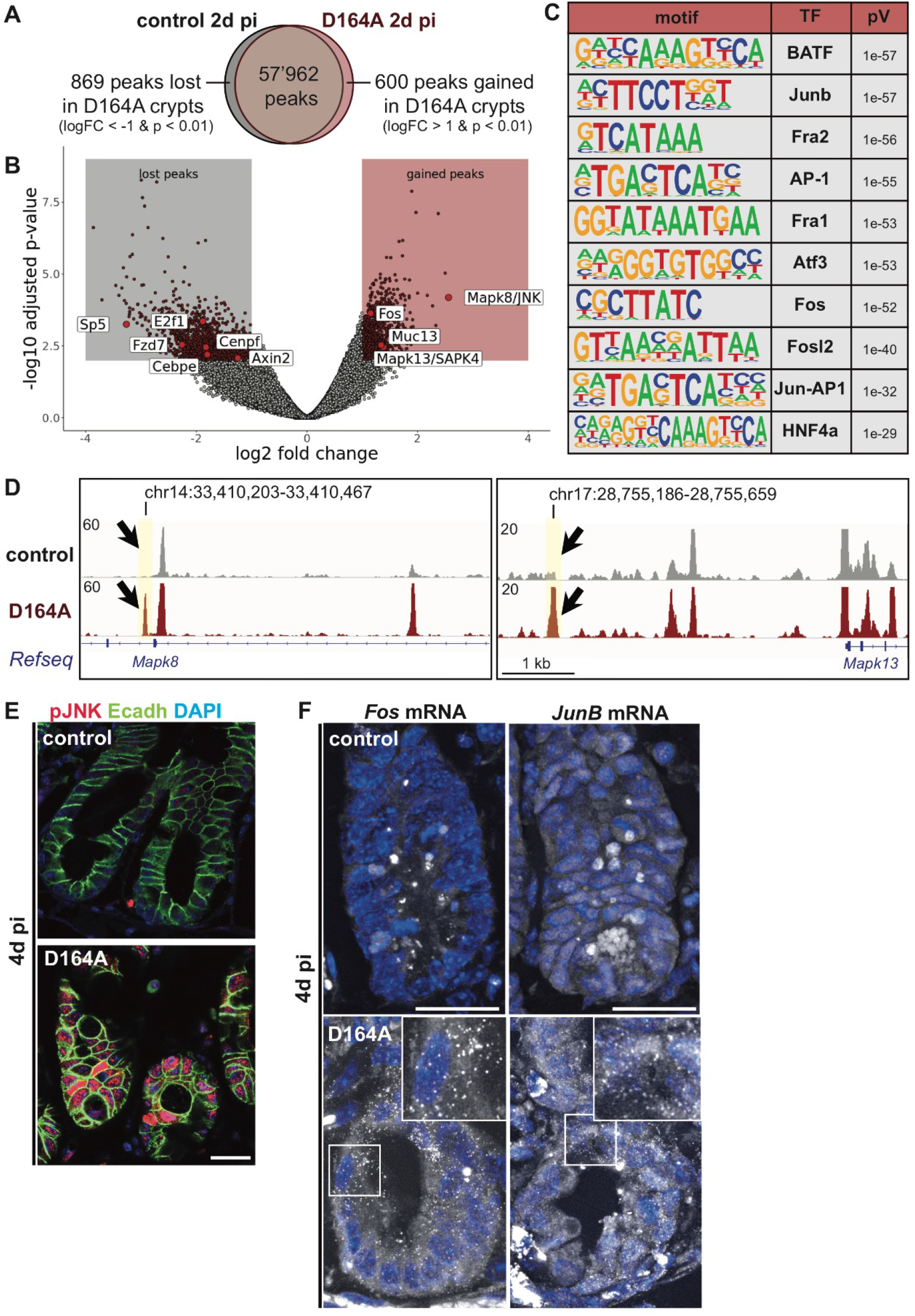
JNK signalling mediates major chromatin remodelling and triggers the expression of villus tip genes in D164A crypts. A-B) Venn diagram and volcano plot of ATACseq peaks in control and D164A crypts. Differential peak analysis indicates that 600 peaks are gained (logFC > 1 & p < 0.01) and 869 are lost (logFC < −1 & p < 0.01) in D164A crypts 2d pi. Selected peaks labeled. C) 10 most significantly enriched transcription factor (TF) binding motifs from HOMER motif analysis on differential peaks. D) Combined ATACSeq tracks of control and D164A counts. Black arrows show gained peaks with AP-1 binding motif annotated to Mapk8/JNK and Mapk13/SAP1. E) Immunostaining of phospho-JNK. E-cadherin (cell shape) and DAPI (nuclei) counterstain. F) smFISH of *Fos* and *JunB* mRNA. Arrows indicate single mRNA molecules visible as dots. Scale bars, 20 μM.

### JNK signalling mediates major chromatin remodelling in D164A crypts

In line with our scRNASeq results, in the ATACSeq data we found a strong enrichment of motifs bound by transcription factors downstream of the c-Jun N-terminal kinases (JNK) signalling, namely the AP-1 transcription factors Fra1/2, Fos, Fosl2, JunB, Atf3 and BATF (Fig. 5c). Cumulatively, these motifs were enriched in 220 out of the 600 gained peaks in D164A (37%), including peaks mapped to *Fos*, as well as *Mapk8* (JNK) and the closely related *Mapk13* (SAPK4), indicating robust JNK-mediated activation of gene expression (Fig. 5d and Extended Data Fig. 4a). We validated JNK pathway activation by immunofluorescence, which reveals strong nuclear accumulation of phosphorylated JNK (pJNK) in D164A, but not in control crypts 4d pi (Fig. 5e). Similar to *Creb3l3*, the expression of transcription factors *Fos* and *JunB* is normally restricted to enterocytes at the villus tip^31,37^. However, in D164A animals 4d pi, *Fos* and *JunB* transcripts are ectopically expressed in crypts (Fig. 5f), corroborating the ATAC-Seq and scRNASeq results. Moreover, the transmembrane mucin glycoprotein *Muc13*, which was independently shown to be induced by JNK activity^38^, also Fig. among the gained peaks in D164A mutant crypts (Fig. 5b), in line with increased alcian blue positivity of hyperplastic and mis-differentiated intermediate cells 4d pi. Altogether, these results indicate that, following lack of β-catenin-NTF interactions, JNK signalling is activated. While mis-differentiation of IESCs upon perturbation of niche signalling has been previously described, this is, to our knowledge, the first time the JNK pathway has been implicated in loss of intestinal epithelial stem cells.

## Discussion

We have previously shown that the C- and N-terminal transcriptional outputs of Wnt/β-catenin signalling have distinct and independent functions during embryonic development^7^. In this study, we aimed to uncover their specific contributions to the maintenance of adult intestinal epithelial stem cell homeostasis. Our data strongly suggest that C-terminally recruited co-activators act as a binary on-off-switch, and are therefore essential for the basal β-catenin-mediated transcription of Wnt-target genes. Conversely, N-terminal co-factors fine-tune β-catenin’s transcriptional output to the levels required for proper proliferation and self-renewal of IESCs. This selective transcriptional modulation may be key in preserving “just-right” levels of Wnt signaling, which have been implicated in IESC homeostasis both physiological and in tumor context^39^.

While increasing attention has been devoted to the role of BCL9/9l in maintaining colorectal cancer stemness, the function of β-catenin-NTF-interactions in healthy intestinal homeostasis has been overlooked. This might be due to fact that BCL9/9L loss in the intestinal epithelium only transiently induces downregulation of IESC markers^12–15^. However, compelling evidence has recently been provided that the intestinal IESC niche is readily regenerated upon perturbations, as differentiated cells can revert to stem cells^40–43^. Epithelial plasticity can thus mask the effects of loss of function studies of proteins, such as BCL9/9L, whose contribution to intestinal homeostasis only becomes apparent when the epithelial stem cell niche is challenged. Our data suggest that N-terminal co-factors selectively restrain Wnt/β-catenin outputs that promote Myc-E2F-driven proliferation, thereby ensuring the maintenance of a self-renewing pool of IESCs. Indeed, in N-terminal mutant (D164A) crypts, excessive proliferation of IESCs rapidly led to exhaustion of the stem cell pool. Concomitantly with robust JNK pathway activation, the crypts underwent a profound mis-differentiation and “villisation”. We show ectopic crypt expression of gene programs characteristic of enterocytes at the villus tip^31,37^, where terminally differentiated cells are eliminated by apoptosis. Mounting inflammation and colitis eventually resulted in a lethal loss of intestinal function. Further studies should dissect the role of immune cell-derived signals for maintenance of intestinal homeostasis in both physiological and perturbed conditions.

The cellular dynamics of crypt collapse in N-terminal mutant animals are in striking contrast with the rapid crypt atrophy observed upon β-catenin deletion, or lack of interactions with C-terminally recruited co-factors. The results presented herein (Fig. 6) provide evidence that discrete regulation of β-catenin transcriptional outputs preserves the narrow window of Wnt-pathway activity and cellular specificity required to govern the identity and fate of intestinal epithelial stem cells.

**Figure 6.**
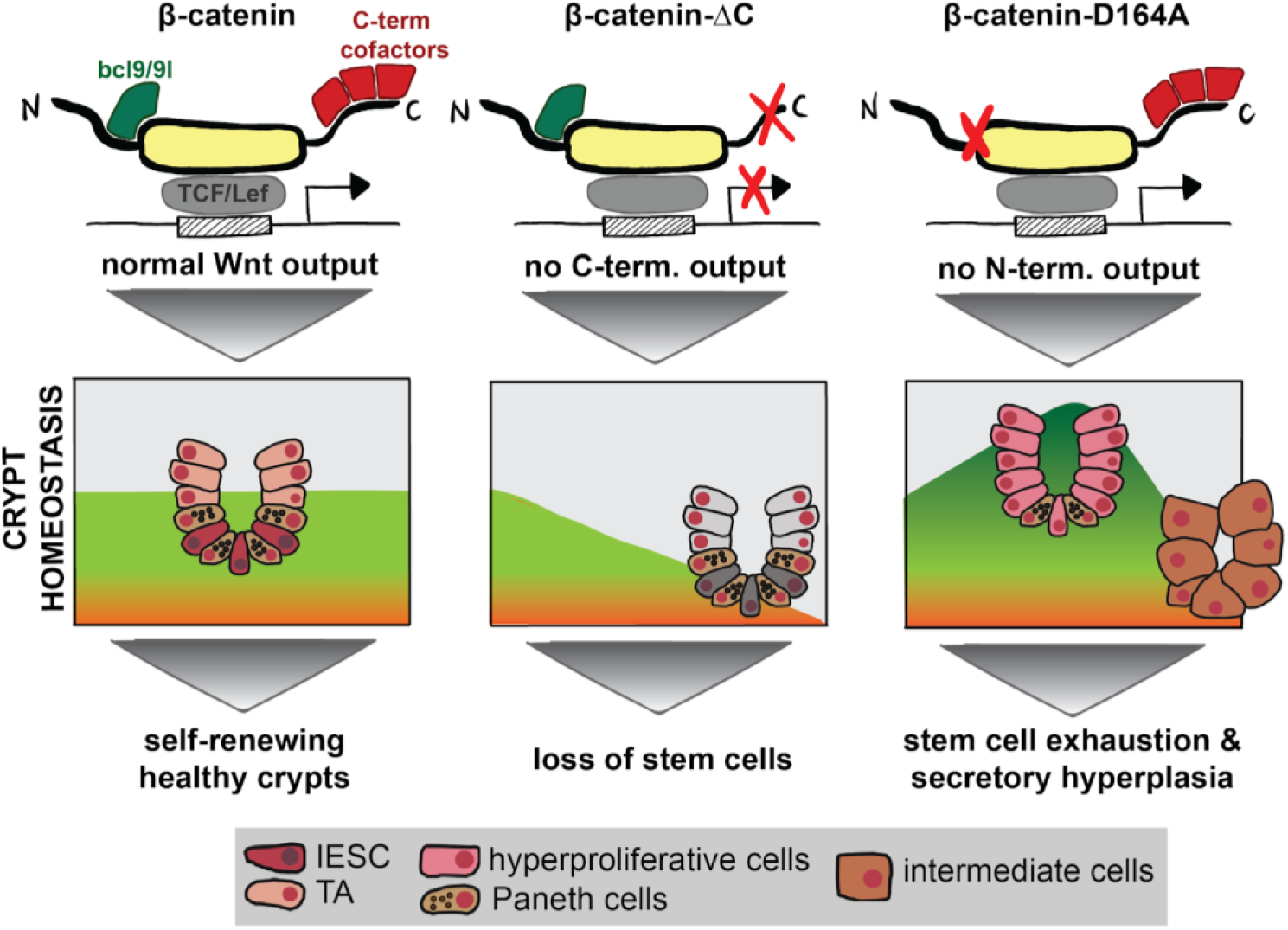
Scheme summarizing the distnct roles of β-catenin transcriptional outputs on the identity of the intestinal epithelial stem cells. C-terminal co-activators govern basal Wnt signaling and preventing their activity leads to loss of stem cells and proliferative arrest. Contrary, N-terminal co-factors act as stem cell specifyers. Blocking their contribution results in hyperproliferation and aberrant differentiation of intestinal epithelial stem cells. (IESC-intestinal epithelial stem cell, TA-transit amplifier cell)

## Author Contributions

CB designed and performed experiments, collected, analyzed and interpreted data and wrote the manuscript. TV initiated and conceived the research, designed and performed experiments, interpreted data and wrote the manuscript. KH, KV and GM assisted with the experiments. KH, AL and LVR helped with the bioinformatic analysis. GH and EAM discussed the data and assisted with manuscript preparation. EAM supported the research. KB initiated and supported the research, discussed the data and assisted with manuscript preparation.

## Acknowledgements

S. Robine kindly shared the *Villin-Cre^ERT2^* strain with us. We are grateful to C. Cantù (Linköping University, Sweden) for valuable comments and to S. Janjuha and W. Macnair for advice. We appreciate discussions with members of the Basler and Moor lab, in particular with N. Doumpas and J. Hilchenbach. For technical help we thank E. Escher and the Functional Genomics Center Zurich. This work was supported by the Swiss National Science Foundation, the Swiss Cancer League, the University of Zurich Research Priority Program (URPP) “Translational Cancer Research” and the Kanton of Zürich. TV is supported by Czech Science Foundation grant 18-21466S and is a fellow of the URPP Translational Cancer Research.

## Conflict of Interetst

The authors declare no conflict of interets.

## Methods

### Mice

The *Ctnnb1-flox* allele^44^ was combined with the *Ctnnb1-delC, Ctnnb1-dm* or *Ctnnb1-D164A*^7^, and bred into the VillinCre-ER^T2^ mouse strain (The Jackson Laboratory) to generate *villin-CreER^T2^;Ctnnb1^wt/flox^* (control) *villin-CreER^T2^;Ctnnb1^flox/flox^* (KO), *villin-CreER^T2^;Ctnnb1^dm/flox^* (dm), *villin-CreER^T2^;Ctnnb1^D164A/flox^* (D164A) and *villin-CreER^T2^; *Ctnnb1*^Δ*C/flox*^* (ΔC). In these mice, the conditional β-catenin allele can be deleted specifically in the intestinal epithelium by tamoxifen-inducible VillinCre-ER^T2^-mediated recombination, leaving Wnt/β-catenin-mediated transcription under the sole control of mutant β-catenin (wt in control animals). To induce Cre-ER^T2^-mediated recombination, tamoxifen (Sigma, 80 mg/kg) was injected intraperitoneally on two consecutive days. Mice were sacrificed 48 hours after the first tamoxifen injection. Alternatively, mice were injected on 4 consecutive days and sacrificed 96 hours after the first injection. Mouse experiments were performed in accordance with Swiss guidelines and approved by the Veterinarian Office of the Kanton of Zurich, Switzerland. All animals were kept on a C57BL/6 background. Mice were 8-12 weeks old at the time of treatments and cell isolations. Mice of both sexes were used in all experiments and littermates were used as controls.

### Duodenal organoids

Duodenal crypts were isolated and cultured as described^45^ with minor modifications. The duodenum was dissected out, flushed thoroughly, opened longitudinally and then cut in 1-2 mm stripes. After repeated washing in ice-cold PBS, the duodenal fragments were incubated for 20 min in Gentle Cell Dissociation Reagent (Stemcell Technologies). Crypts were gradually released from the tissue during 4 rounds of washing in 0.2% BSA and filtering through a 70 micron filter, generating 4 fractions. Only fractions 3 and 4 containing intestinal crypts were used for downstream experiments (organoids or single cell suspension). Crypts were seeded in 50 μL Matrigel (Corning) domes and cultured in Intesticult (Stemcell Technologies). Upon passaging, the culture medium was supplemented with 1 μM Rho-associated protein kinase (ROCK) inhibitor (Y27632, Millipore). Recombination was induced with 100 nM 4-hydroxytamoxifen (4OHT, Sigma) added to the culture medium upon splitting.

## METHOD DETAILS

### Mouse genotyping

The *Ctnnb1-D164A* allele was detected with the primers 5’-TCCCTGAGACGCTAGATG-3’ and 5’-GAGTCCCAGCAGTACAAC-3’, yielding an amplicon of the size 475 bp for the wild-type and 628 bp for mutant alleles^7^. *Ctnnb1-ΔC* allele was determined using primers 5’-GTGCACACGTCATGCTTTAC-3’ and 5’-TGGCTTGTCCTCAGACATTCG-3’, which generate an amplicon of size 349bp for the wild-type and 415 bp for mutant alleles^7^. The presence of the *villin-CreER^T2^* allele was detected with the primers 5’-CAAGCCTGGCTCGACGGCC-3’ and 5’-CGCGAACATCTTCAGGTTCT-3’, which generate a 220 bp product^16^. Recombination of the conditional β-catenin allele was confirmed for every mouse included in this study by PCR with the primers 5’-AAGGTAGAGTGATGAAAGTTGTT-3’ (RM41), 5’-CACCATGTCCTCTGTCTATTC-3’ (RM42) and 5’-TACACTATTGAATCACAGGGACTT-3’ (RM43), generating products of 221 bp for the wild-type allele, 324 bp for the floxed allele and 500 bp for the floxdel allele^44^ (Extended data Fig. 1d).

### Immunohistochemistry

Dissected duodenum was flushed thoroughly in ice-cold PBS, then cut in 1 cm pieces and fixed in 4% PFA in PBS overnight at 4°C. After repeated washing in PBS, tissues were dehydrated in a spin tissue processor, embedded in paraffin and cut in 8 μm sections. Deparaffinized tissue sections were subjected to antigen retrieval in 2.4 mM sodium citrate and 1.6 mM citric acid, pH 6, for 25 min in a steamer. Sections were washed with PBST (0.1% Tween-20 in PBS) and blocked for 30 min at RT in blocking buffer (5% BSA, 5% heat-inactivated normal goat serum in PBST). Following overnight incubation at 4°C with primary antibody (1:100, in blocking buffer), sections were washed in PBST and incubated with secondary antibody (1:400, in blocking buffer) for 1 hr at room temperature. Nuclei were stained with DAPI (Sigma, 1:1000) in blocking solution for 5 min at RT. Sections were imaged on a Leica LSM 710 confocal microscope and processed equally using the ImageJ (FIJI) software, or scanned on Vectra 3.0 Automated Quantitative Pathology Imaging System (Perkin Elmer).

### Double histology for intermediate cells

Deparaffinized tissue sections were incubated with hydrogen peroxide for 5 min to block endogenous peroxidase, then incubated for 30 min in blocking buffer (5% BSA, 5% heat-inactivated normal goat serum in PBST) for 30 min. After overnight incubation with primary anti-Lyz antibody (Dako), sections were repeatedly washed in PBST and incubated with biotinylated secondary antibody for 1h at RT and then visualized with VECTASTAIN ABC HRP Kit (Vector Laboratories) as indicated by the manufacturer. Sections were then incubated in a 3% aqueous solution of acetic acid for 3 min and then mucins were stained with alcian blue (1g in 100 mL 3% acetic acid) for 15 min. Hematoxylin (50% aqueous solution) and eosin staining were then carried out according to standard procedure. Sections were scanned on Vectra 3.0 Automated Quantitative Pathology Imaging System (Perkin Elmer).

### RNA *in situ* hybridization

Duodenum was flushed thoroughly in ice-cold PBS, then cut in 1 cm pieces and fixed under gentle agitation in 10% formalin for 20 hrs at room temperature. *Axin2* mRNA *in situ* hybridization was performed with the ACD RNAscope kit (ACDBio), according to the manufacturer’s instructions. Sections were image on a Vectra 3.0 Automated Quantitative Pathology Imaging System and quantification was performed with the inForm Cell Analysis software (Perkin Elmer).

### Single molecule *in situ* hybridization

Mice were sacrificed and the duodenum was removed and flushed thoroughly in ice-cold PBS. Duodenal tissue was then cut in 1 cm pieces and fixed in 4% PFA (Santa Cruz Biotechnology, sc-281692) in PBS for 3 hours and subsequently agitated in 30% sucrose, 4% PFA in PBS overnight at 4°C. Fixed tissues were embedded in TIssue-Tek OCT Compound (Sakura, 4583). 8 μm thick sections were sectioned onto poly L-lysine coated coverslips. Probe libraries were designed using the Stellaris FISH Probe Designer Software (Biosearch Technologies) (see Extended data Table 1) and coupled to Cy5 (*Myc, Fos, Creb3l3*) or TMR (*Lgr5, JunB)* as described^46^. The intestinal sections were hybridized with smFISH probe sets according to a previously published protocol^47^. DAPI (Sigma-Aldrich) was used as nuclear counterstain. smFISH imaging was performed on a Leica THUNDER 3D Live Cell Imaging system using the following THUNDER Computational Clearing Settings, Feature Scale (nm): 350, Strength (%): 98, Deconvolution settings: Auto and Optimization: High.

### Organoid formation assay and EdU staining

Mice were sacrificed 0, 2 or 4d post injection, crypts were isolated as described^45^. Fractions 3 and 4 were seeded in equal amounts (500 crypts) in 50 μL Matrigel domes. Organoids were quantified manually 7 day post crypt seeding and micrographs were collected using the AmScope 5.2. software. For EdU staining, crypts were seeded on glass-bottomed 8-well chambered slides (Thermo Fisher Scientific, Lab-TekTM, 154532) in 20μL drops. EdU incorporation (30 min pulse) and visualization was performed according to the manufacturer instruction with Edu Click 647 Kit (baseclick) prior to overnight incubation with anti-Ecadherin (BS Transduction Lab) antibody for cell shape staining. Imaging was performed on a Visitron CSU-W1 spinning disk confocal microscope with a CFI PlanFluor 20x objective.

### RNA sequencing

Control (n=3), KO (n=2), dm (n=3), ΔC (n=2) and D164A (n=3) mice were sacrificed 2d pi and the duodenal epithelial RNA isolation was performed as described^48^. Briefly, dissected duodenum was flushed thoroughly in ice-cold PBS, then opened longitudinally and incubated for 20 min on ice in dissociation buffer 1 (30 mM EDTA, 1.5 mM DTT in PBS), followed by 10 min incubation in dissociation buffer 2 (30 mM EDTA in PBS) at 37 °C under gentle agitation. The epithelium was released by vortexing, pelleted by centrifugation at 4 °C for 5 min at 1000g, and resuspended in 500 μL TRI Reagent (Sigma). RNA was extracted by phenol-chloroform precipitation. DNAse treatment was carried out with DNA-free DNA Removal Kit (Invitrogen) according to the manufacturer’s instructions. Library preparation was performed with the Illumina TruSeq RNA Kit. RNA sequencing was performed on the Illumina NextSeq500 by the Functional Genomics Centre Zurich (FGCZ).

### Droplet-based single cell mRNA sequencing

Control (n=3) and D164A (n=3) mice were sacrificed 0, 2 or 4d post injection, crypts were isolated as described^45^. Crypt fractions 3 and 4 were pooled and dissociated to single cells by incubating them with 5 mL pre-warmed TripLE Express Enzyme (ThermoFischer) in a 37 °C water bath for 5 min with frequent agitation. TripLE was inactivated with 50% FBS in Advanced DMEM/F12 (ThermoFischer). Single cells were pelleted at 180 g for 3 min and resuspended in 2 mL cold Intesticult (Stemcell Technologies Inc). Clumps were dissolved by pipetting up and down with a 1mL pipet before filtering twice through a 40 micron filter. Cell viability and number were quantified in automatically with a Countess (Thermo Scientific). DropSeq workflow was performed as described^49^ on a Nadia Instrument (Dolomite Bio). Sequencing was performed on the Illumina NovaSeq6000 with an SP Reagent Kit.

### ATAC-Seq

Control (n=3) and D164A (n=3) mice were sacrificed 2d pi and processed independently for ATACSeq library preparation, following the protocol by^50^. Briefly, single cell suspension of duodenal crypts was performed as for scRNAseq, which ensures virtually absent contamination from non-epithelial cells. 50’000 cells were lysed in 50ml cold Lysis buffer (10 mM Tris-HCl,pH 7.4, 10 mM NaCl, 3 mM MgCl2, 0.1% IGEPAL CA-630). After centrifugation at 500 g for 10 min at 4°C, cells were resuspended in 50 mL 1X Illumina transposition mix (25ml Tagment DNA Buffer, 2.5ml Tagment DNA Enzyme, 22.5ml nuclease-free H2O) and incubated for 30 min at 37°C on a shaker. Immediately following the transposition reaction, DNA was purified using the QIAgen MinElute PCR Purification Kit according to the manufacturer’s instructions. Library was amplified with Nextera Sequencing primers in NEBNext Hot Start High-Fidelity 2X PCR master mix for 5 cycles. The appropriate number of additional PCR cycles was determined by qPCR. Amplified libraries were purified using the QIAgen MinElute PCR Purification Kit. DNA was eluted in 20ml EB buffer and quantified and visualized for quality control with a 2200 TapeStation System (Agilent). Libraries were sequenced on an Illumina HiSeq2500 with paired end 70 bp read configuration.

## QUANTIFICATION AND STATISTICAL ANALYSIS

### Quantification of immunofluorescence and RNA *in situ*

Percentage of crypt cells positive for Ki67 antibody staining and Axin2+ RNA *in situ* hybridization were automatically quantified with the software inForm Cell Analysis software (Perkin Elmer). Crypt area was manually defined. We quantified at least 30 crypts from 2-3 different mice per condition. Barplots were generated on GraphPad Prism. The unpaired Student’s T test function in GraphPad Prism was used to analyze the significance of two-group comparisons.

### smFISH quantification

The of Lgr5+ area per crypt was manually quantified using FIJI. We quantified at least 8 crypts from 3 different mice per condition. Barplots were generated on GraphPad Prism. The unpaired Student’s T test function in GraphPad Prism was used to analyze the significance of two-group comparisons.

### Bulk RNA-seq analysis

Reads were quality-checked with FastQC. Reads alignment to the reference genome “Mus_musculus.GRCm38.95” and read count was performed on the Support Users for SHell script Integration (SUSHI) framework^51^, with the RSEMApp application. Pairwise comparisons were performed with the SUSHI application EdgeRApp (based on edgeR^52^). Principal component analysis and filtering of differentially expressed genes (DEGS, logFC > |2|, p < 0.01) were performed on R (version 3.6.1). The package pheatmap^53^ was used to generate the heatmap of normalized FPKM. We performed gene set enrichment analysis^18^ on significantly differentially expressed genes (logFC > |2|, p < 0.01) using the Bioconductor package fgsea with default parameters^54^. Genes were ranked based on p-value, and taking into account directionality of the fold change with the formula *ranking* = −*log*10(*P*)/*sign*(*log*2*ratio*) (obtained from the blogpost http://genomespot.blogspot.co.at/2016/04/how-to-generate-rank-file-from-gene.html). The Hallmarks gene set collection from the Molecular Signatures Database^55^ was imported in R with the package msigdbr. Cell type enrichment analysis of DEGs across conditions was performed on the web-based tool EnrichR^56,57^.

### Computational analysis of scRNaseq data

#### Dimensionality reduction and clustering

Reads were quality-checked with FastQC. Raw data processing of fastq files was performed with the zUMIs pipeline^58^ using the reference genome “Mus_musculus.GRCm38.95". scRNaseq analysis was performed with Seurat 3 on R 3.6.1^59^. Cells were filtered based on mitochondrial gene content, unique molecular identifier (UMI) counts were log-normalized according to default Seurat settings. Scaling was performed on the variable genes (FindVariableFeatures, parameters: x.low.cutoff = 0.0125, x.high.cutoff = 3, y.cutoff = 0.5) regressing out UMI number and mitochondrial gene content. Samples were merged and joint analysis was performed with the package conos^60^. Cells were clustered with the leiden community method (resolution=1.3). The joint graph (in PCA space) was embedded in UMAP space and converted to a Seurat object. The stem cell and early progenitor clusters were subsetted and the log-normalized UMI counts were exported and used for the normalization algorithm and downstream analysis.

#### Normalization algorithm

Analysis of the control timecourse data revealed that the acute loss of one β-catenin allele only caused a slight and transient downregulation of Wnt-targets, proliferation and stem markers, which then returned to normal expression levels by 4d pi. Thus, we devised a normalization of the mutant timecourse data to disentangle these effects from those induced by impaired N-terminal interactions, which were the focus of our investigation. For every timepoint (0, 2 and 4d pi), we divided the D164A single-cell expression (from data slot) of each gene by the mean expression of the corresponding gene in the control with the formula: *ln*[*exp*(*D*164*A*)/(*mean*(*exp*(*control*)]. The resulting three normalized D164A sparse matrices (one per timepoint) were used to create Seurat objects, which were merged, scaled and visualized in UMAP space using dimensions 1:10.

#### Diffusion maps and pseudotime analysis

For diffusion maps and supervised pseudotime analysis, the normalized Seurat object was converted in a SingleCellExperiment. Diffusion maps were generated with the Bioconductor package destiny^61^ using default parameters. Scores in the diffusion component 1 were compared using the Wilcoxon test (alternative: two.sided) in R base. Supervised pseudotime analysis was performed with the Bioconductor package psupertime^26^ on all genes or on mouse transcription factors only, setting scale to FALSE.

#### Cell cycle scoring and classification

Cell cycle scoring was performed with the CellCycleScoring algorithm from Seurat, using cell cycle-related genes from^25^. Wilcoxon test (alternative: two.sided) was used to compare G2M scores. To assign a measure of similarity to IESCs or TAs transcriptomic profiles, we trained a logistic regression (modification of the multinomial regression of MatchSCore2^62^ using IESC and TA markers extracted from a publicly available single cell dataset^29^. Upon collecting the estimated probabilities of TA class memberships for each timepoint separately, we compared these probability distributions and tested the significance of the difference using Kolmogorov-Smirnov test (alternative: two.sided).

#### Gene set enrichment analysis

We used the Seurat FindMarkers function (default parameters min.pct 0.25 and logfc.threshold 0.25) to compute differentially expressed genes between timepoints in our normalized D164A stem cell and early progenitor dataset. We then ranked genes with the formula *ranking* = −*log*10(*p_val_adj*)/*sign*(*avg_logFC*) and performed gene set enrichment analysis^18,55^ on R using the Bioconductor package fgsea with default parameters^54^. The following gene set collections were imported in R with the package msigdbr: Hallmarks^55^, Chemical and Genetic Perturbations (various contributions), KEGG^63^, ^64^and gene ontology biological process^65^. Plots were generated with the R package ggplot2^66^.

#### Computational ATAC-Seq analysis

Reads were quality-checked with FastQC. Adapters trimming with cutadapt, alignment on GRCh38.95 using Bowtie2 with default mapping parameters^67^, as well as duplicate filtering with Picard and peak calling with MACS2 (p<0.01) were performed on the ENCODE pipeline^68^ supported by the SUSHI application AtacENCODEApp, for every biological replicate separately. The peak files were merged in BEDtools v2.29.2^69^ and converted into a saf annotation, to which raw reads were mapped with the FeatureCounts function in the package rsubread^70^. The resulting count matrix was filtered for peaks with low coverage (minimum read count of 10 for each sample). The remaining counts were then normalized by total library size and TMM-derived normapllization factors calculated with edgeR^52^. Differential peaks between sample groups were identified using the exact test function in edgeR. Only peaks with a log fold change > 1 and an p value < 0.01 were considered as differentially accessible for further analysis. HOMER^36^ was used for peak annotation and motif enrichment analysis. Motifs with Benjamini-corrected q-value < 0.001 were considered significantly enriched. bigWig files of control (n=3) and D164A (n=3) were combined into a single track for visualization in Integrative Genomics Viewer^71^. Genes associated with a differentially accessible peak were ranked with the formula *ranking* = −*log*10(*pvalue*)/*sign*(*log2FC*) and used for gene set enrichment analysis with the R using the Bioconductor package fgsea with default parameters^54^. Significance of overlap between gene sets overrepresented in scRNAseq and in ATACSeq was quantified with hypergeometric test. Plots were generated with the R package ggplot2^66^.

## DATA AND CODE AVAILABILITY

Data generated in this study are deposited in GEO with the accession numbers: GSE148941, GSE148942, GSE148940. The code used in this study are available at the repository https://github.com/cocoborrelli/betacat.

## Extended Data

### Extended Data Figure Legend

**Extended Data Figure 1.**
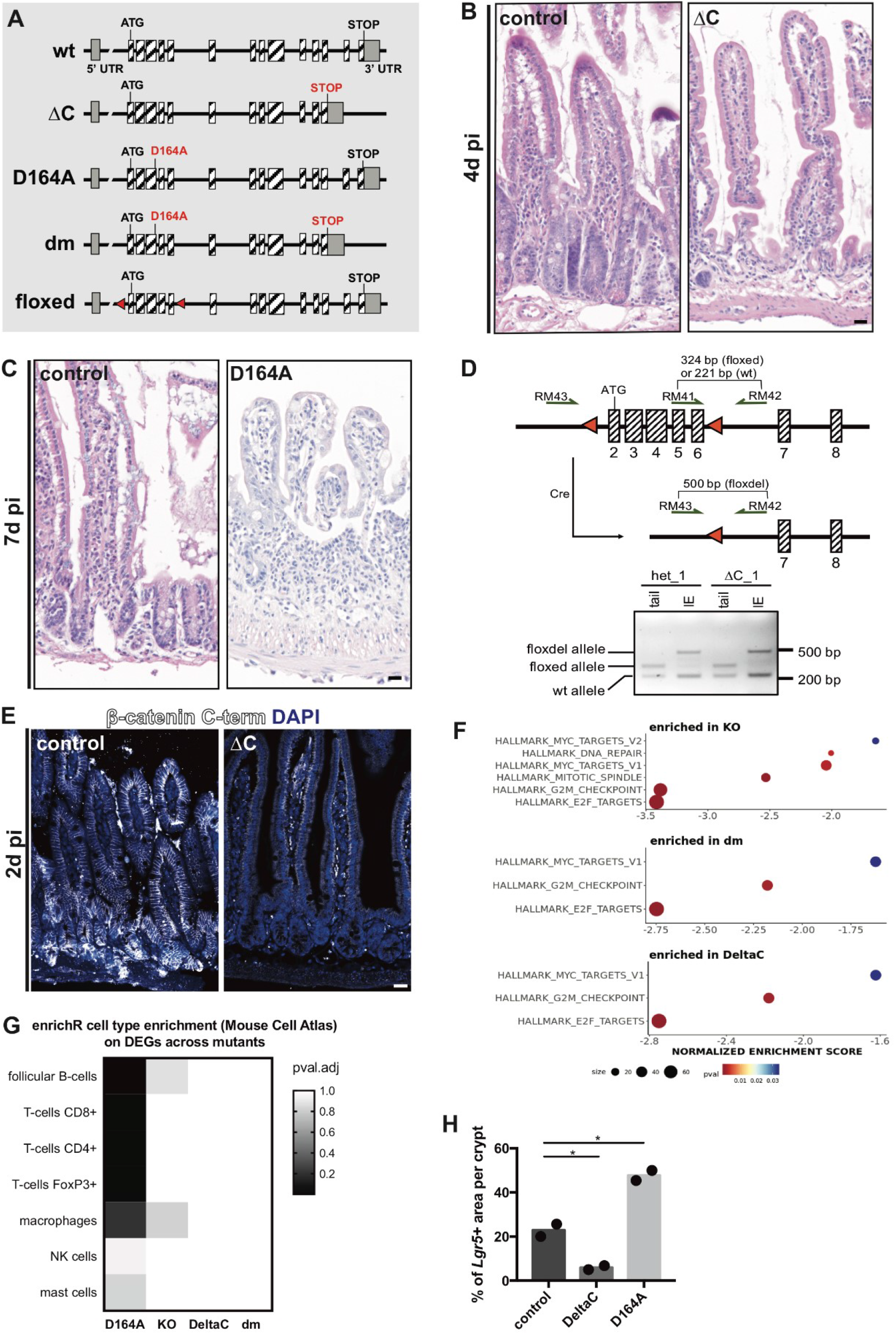
Distinct effects of N- vs. C-terminal β-catenin transcriptional outputs on intestinal homeostasis. A) Scheme of wt, mutant and conditional *Ctnnb1* alleles used in this study. B) ΔC animals suffer from crypt atrophy and reach humane endpoint 4d pi. C) D164A (N-terminal mutant) animals show loss of crypts and villus shortening 7d pi. Thickening of the mesenchyme indicates immune infiltration and severe colitis. D) Map of the β-catenin locus, modified from^44^ depicting how the recombination of conditional allele was confirmed. Red triangles indicate the loxP sequences flanking exons 2 (which contains the ATG) to 6. Prior to recombination, primers RM41 and RM42 generate a 221 bp band for the wt allele and a 324 bp band for the floxed allele, respectively. Upon Cre induction, primers RM42 and RM43 generate a 500 bp band. Representative PCR showing successful and tissue-specific recombination in DNA obtained from the intestinal epithelium but not from the tail. E) Depletion of wt (flox) β-catenin in the crypts of ΔC animals 2d pi, as shown by immunofluorescence with antibody specific for β-catenin’s C-terminal moiety. F) Gene set enrichment analysis (GSEA) on differentially expressed genes (DEGs, logFC > |2|, p < 0.05) of KO, dm and ΔC animals. Annotated gene sets obtained from the Hallmarks collection of the Molecular Signatures Database (MSigDB). G) Cell type enrichment analysis of DEGs across mutants performed on the web-based tool EnrichR^56,57^. H) Percentage of Lgr5+ (IESC marker) area per crypt in control and D164A animals. Student’s T-test, unpaired, * p<0.05. Scale bars, 20 μM.

**Extended Data Figure 2.**
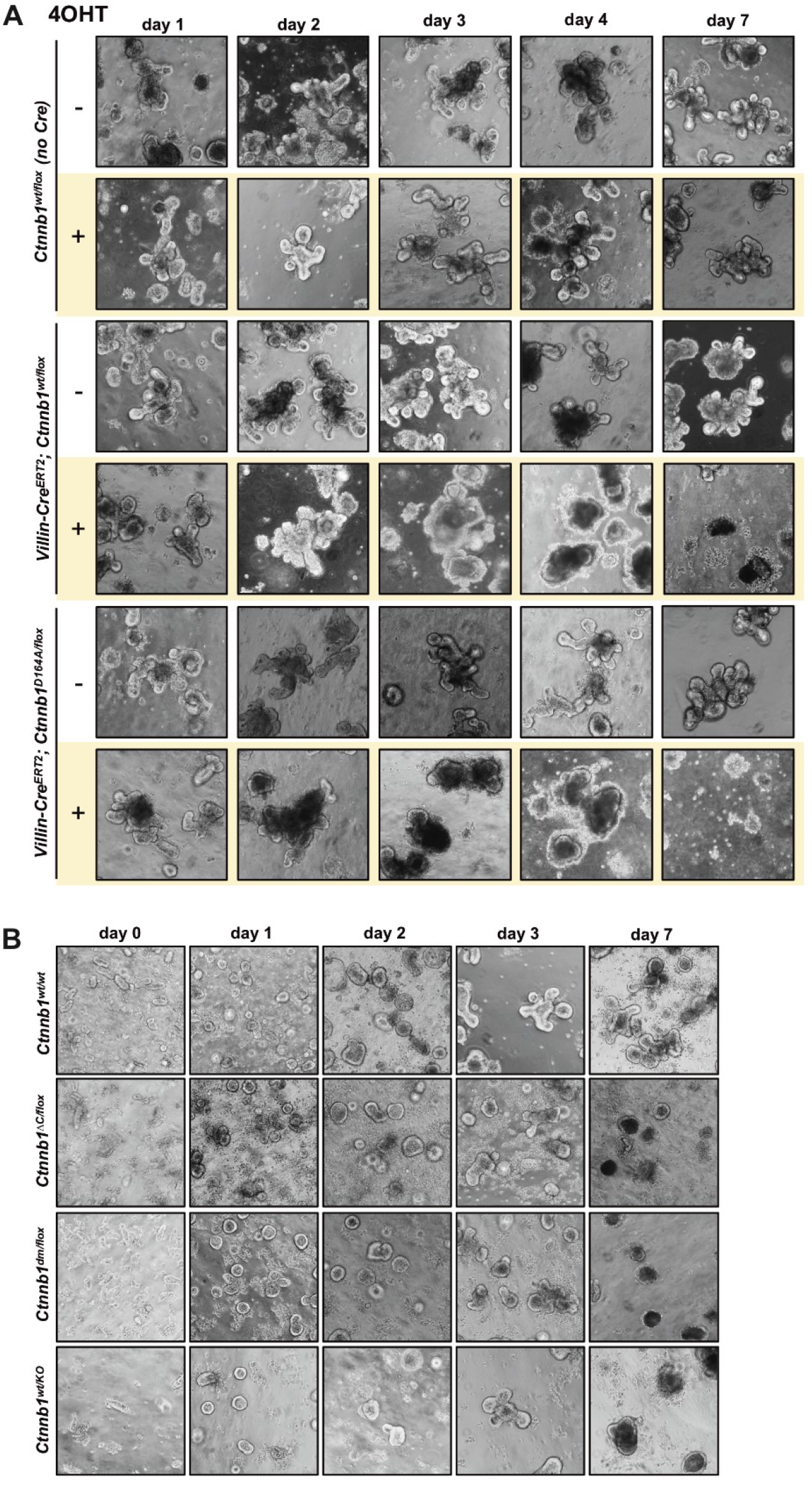
β-catenin is haploinsufficient *in vitro*. A) *villin-CreER^T2^;Ctnnb1^wt/flox^* and *villin-CreER^T2^;Ctnnb1^D164A/flox^* organoids treated with 4-hydroxytamoxifen (4OHT) *in vitro* die 7 days after recombination. Duodenal organoids lacking the *villin-CreER^T2^* allele are not affected by 4OHT. Days after addition of 4OHT indicated at the top. B) Duodenal organoids derived from *Ctnnb1^ΔC/flox^* and *Ctnnb1^dm/flox^*, as well as from constitutively hemizygous *Ctnnb1^KO/wt^* animals mice grow slower than wt controls and can not be propagated *in vitro*. 20x magnification.

**Extended Data Figure 3.**
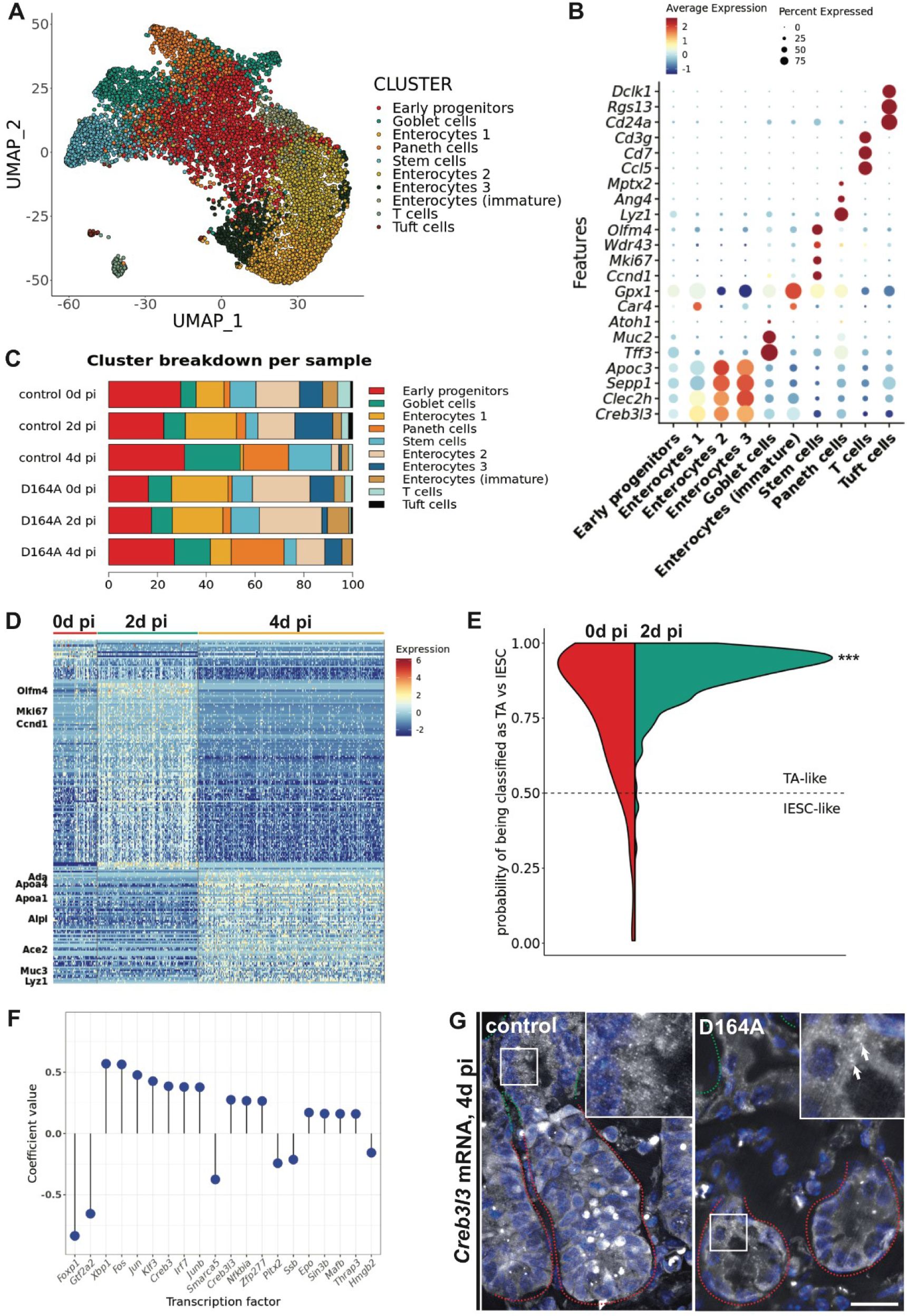
Longitudinal scRNASeq of control and N-terminal mutant (D164A) crypts. A) UMAP embedding of joint graph of single cells sequenced from control and D164A crypts isolated 0, 2 and 4d pi. Leiden clustering results in 10 clusters corresponding to the main cell populations in the intestinal crypt. B) Marker gene expression across manually annotated clusters. C) Cluster breakdown in percent across all samples. D) Heatmap of the top 200 differentially regulated genes in normalized D164A stem cells and early progenitors (SCEP) cells across timepoints. IESC and proliferation markers and upregulated 2d pi, while expression of differentiation markers increases 4d pi. E) Distribution of predicted model responses of logistic regression showing a shift of SCEPs 2d pi towards TA traits. Kolmogorov-Smirnov test (*** p < 0.001). F) Supervised pseudotime ordering coefficients of mouse transcription factors in decreasing absolute value. G) smFISH of *Creb3l3* mRNA evidences its ectopic expression in D164A crypts 4d pi. Red and green dotted lines indicate crypt and villus area, respectively. Scale bars, 20 μM.

**Extended data Figure 4.**
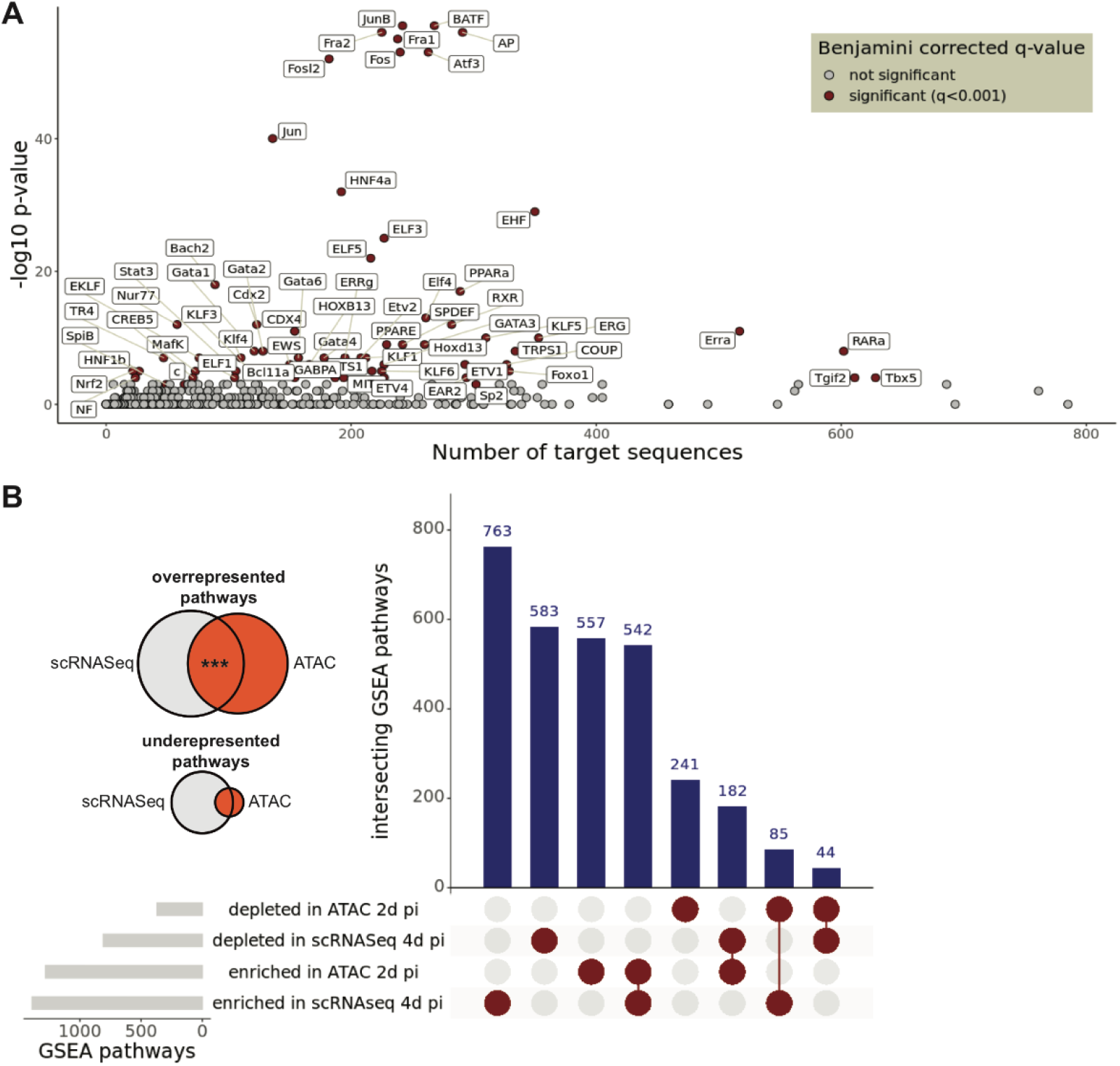
ATAC sequencing of control and N-terminal mutant (D164A) crypts. A) Results of HOMER motif enrichment on 1469 differential ATAC peaks (logFC > |1| & p < 0.01) between control and N-terminal mutant (D164A) crypts. Number of target sequences on x-axis, −log10(p-value) of enrichment on y-axis. Significantly enriched motifs (Benjamini-corrected q-value < 0.001) depicted in red and labeled with corresponding transcription factors. B) Venn diagrams and upset plot showing overlaps between enriched pathways (logFC > |1|) in N-terminal mutant (D164A) crypts, compared to controls, as revealed by ATACSeq (2d pi) and scRNASeq (4d pi). Significance of the overlap of overrepresented pathways calculated with hypergeometric test (*** p < 0.001).

### Extended Data Methods

#### KEY RESOURCES TABLE

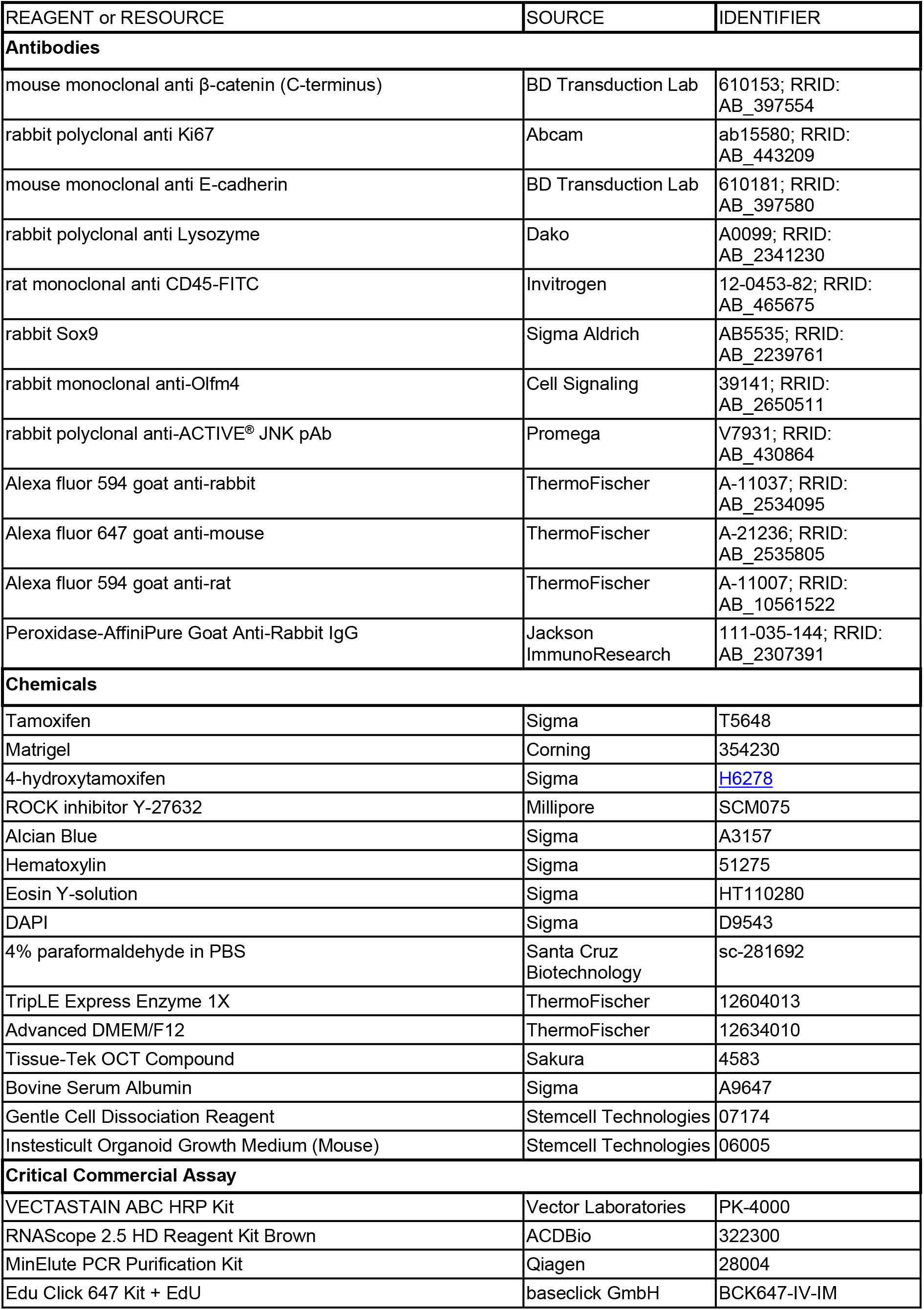
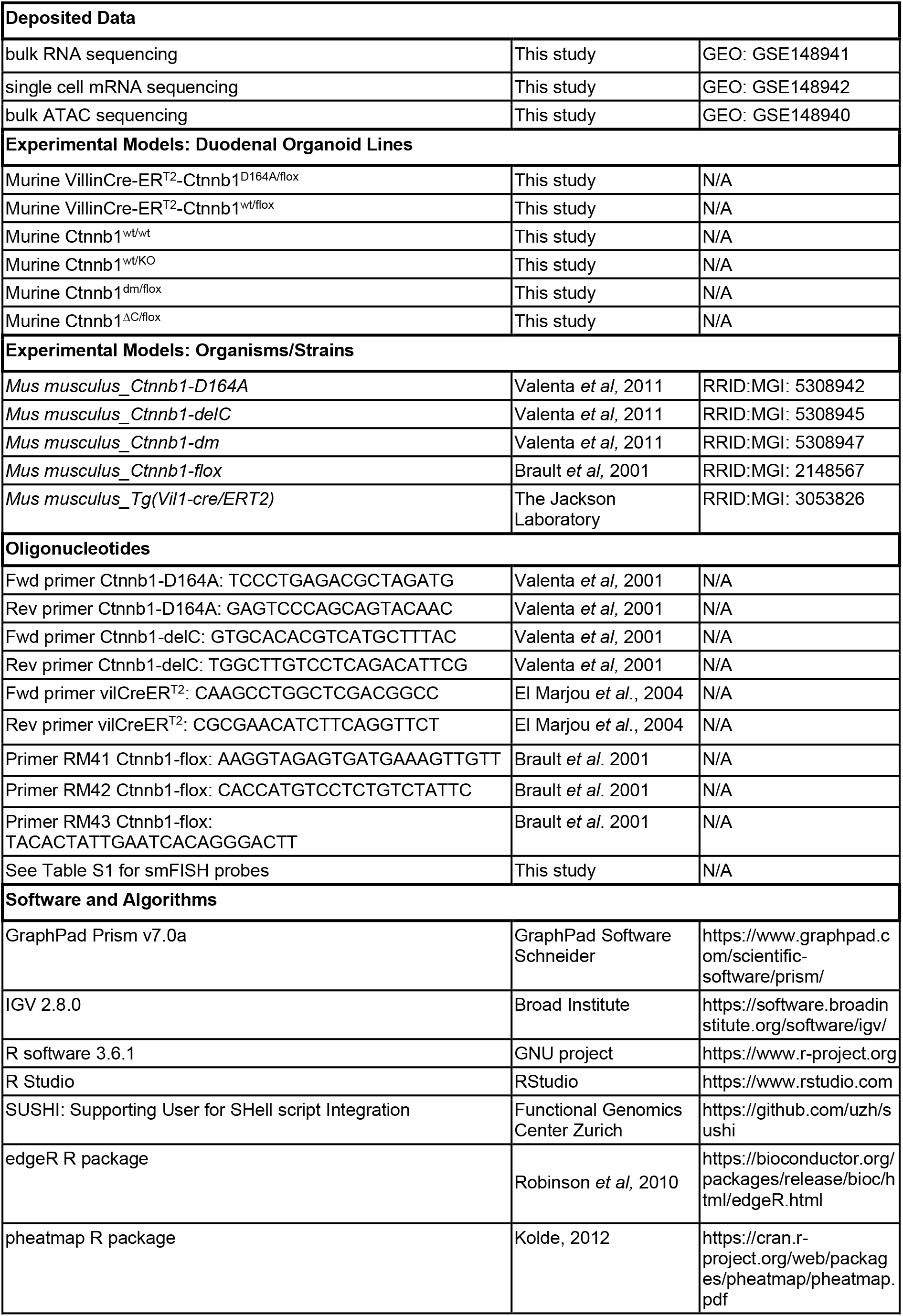
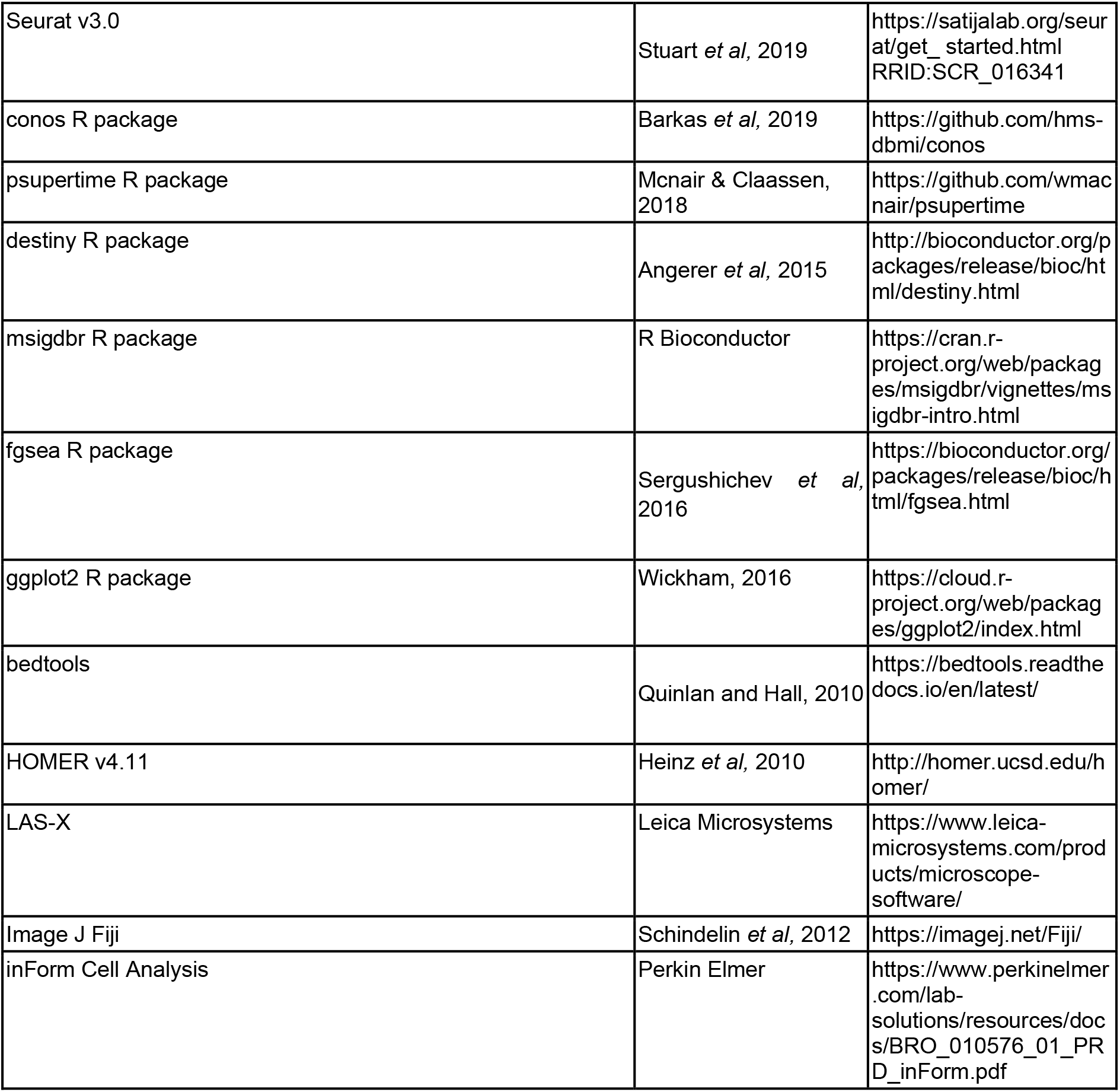

